# The long non-coding RNA *H19* drives the proliferation of diffuse intrinsic pontine glioma with H3K27 mutation

**DOI:** 10.1101/2021.07.19.452516

**Authors:** David Roig-Carles, Holly Jackson, Katie Loveson, Alan Mackay, Rebecca Mather, Ella Waters, Massimiliano Manzo, Ilaria Alborelli, Jon Golding, Chris Jones, Helen L. Fillmore, Francesco Crea

## Abstract

Diffuse intrinsic pontine glioma (DIPG) is an incurable paediatric malignancy. Identifying molecular drivers of DIPG progression is of utmost importance. Long non-coding RNAs (lncRNAs) represent a large family of disease- and tissue-specific transcripts, whose functions have not been yet elucidated in DIPG. Here, we study the oncogenic role of the development-associated *H19* lncRNA in DIPG. Bioinformatic analyses of clinical datasets were used to measure the expression of *H19* lncRNA in paediatric high-grade gliomas (pedHGG). Expression and sub-cellular location of *H19* lncRNA was validated in DIPG cell lines. Locked nucleic acid antisense oligonucleotides were designed to test the function of *H19* in DIPG cells. We found that *H19* expression was higher in DIPG vs normal brain tissue and other pedHGGs. *H19* knockdown resulted in decreased cell proliferation and survival in DIPG cells. Mechanistically, *H19* buffers *let-7* microRNAs, resulting in up-regulation of oncogenic let-7 target (e.g *SULF2*, *OSMR*). *H19* is the first functionally characterized lncRNA in DIPG and a promising therapeutic candidate to treat this incurable cancer.

## INTRODUCTION

Malignant brain tumours are a leading cause of death in paediatric patients. Diffuse intrinsic pontine glioma (DIPG) is a type of paediatric high grade glioma (pedHGG) originating in the brainstem and affecting children with a median age of 6 to 7 years (1,2). DIPGs are highly infiltrative and belong to the fibrillary astrocytoma family where lesions are classified either WHO Grade III or IV (3). At the molecular level, substitution of the lysine at the 27 position of the histone 3 *locus* with methionine (H3K27M at either *H3.1* or *H3.3*.) has been suggested to drive the oncogenesis of DIPGs (4). The H3K27M substitution is identified in approximately 80% of histologically confirmed DIPGs. The World Health Organization has recently classified DIPGs in the broader category of diffuse midline gliomas (DMG) with H3K27M mutation (2,5–7). Another frequent mutation of the *H3 locus*, GR4R is more prevalent on pedHGGs located in the cerebral hemispheres (8). The surgical removal of DIPGs is almost impossible due to their infiltrative nature and to their location. Currently, radiotherapy is the standard treatment for this malignancy; despite this treatment, the median survival for DIPG patients is 9-11 months (9,10). Therefore, novel therapeutic targets are necessary to improve the prognosis of these paediatric patients.

There are approximately 60,000 long non-coding RNAs (lncRNAs) in the human genome; these transcripts are defined as non-coding transcripts longer than 200 nucleotides. Some lncRNAs are highly expressed under pathological conditions and regulate key aspects of tumour progression, invasion and metastasis (11,12). Previous studies have identified several lncRNAs associated with DIPG progression (13,14). However, no lncRNA has been functionally characterised in this malignancy.

*H19* (Gene ID: 283120) is an extensively studied lncRNA, whose aberrant expression has been linked to alterations during foetal development (15). The oncogenic role of *H19* has been described in several malignancies, including adult gliomas (16–18). Mechanistically, *H19* expression is triggered by hypoxia; this lncRNA can affect a wide range of hub genes including microRNAs (miRNAs) and mRNAs (19,20). However, the function of *H19* in pedHGG still remains largely unknown.

Here, we have investigated the clinical relevance of *H19* in DIPG cells and tested *H19*-targeting for halting DIPG proliferation.

## RESULTS

### H19 is up-regulated in DIPG tissue

To confirm the expression of *H19* lncRNA in DIPG tissue, we analysed open-access clinical datasets of pedHGGs and normal brain tissue. Bioinformatic analyses confirmed that *H19* levels were significantly increased in DIPG tissue on “Allis - 45 - custom - ilmnht12v4” dataset (*P*<0.01, Figure 1A) whereas an increasing trend was observed in “Paugh - 37 - MAS5.0 - u133p2” (Figure 1B). Microarray data from Paediatric Cbioportal (https://pedcbioportal.kidsfirstdrc.org/) demonstrated significantly higher *H19* expression in brainstem than hemispheric pedHGGs (*P*<0.01). (Figure 1C). We then explored the influence of mutations in *H3* genes (H3K27M and H3G34R) on *H19* expression; this analysis revealed that H3K27M-bearing pedHGGs expressed higher levels of *H19*, compared to pedHGG bearing H3G34R mutation (*P*<0.0001) or the wild-type *H3* gene (*P*<0.01) (Figure 1D). Histological classification of pedHGG samples showed that DIPG tissues expressed higher *H19* levels than other pedHGGs, including anaplastic astrocytoma and glioblastoma (*P*<0.01) (Figure 1E). Overall survival analysis did not identify significant differences between high- and low-*H19* expressing DIPG groups in CBioportal dataset (median survival was 9.8 and 11.1 months, respectively) (Figure 1F). Analysis of a single-cell RNA-Sequencing dataset from the developing human midbrain showed that *H19* is most strongly expressed by oligodendrocyte precursor cells (OPCs), which are thought to be the cell of origin of DIPGs (21,22) (Figure 1G). All these results indicate that *H19* is highly and selectively up-regulated in DIPGs with H3K27M mutation. Hence, we decided to study the cellular function and mechanism of action of this lncRNA in DIPG-H3K27M.

**Figure 1.**
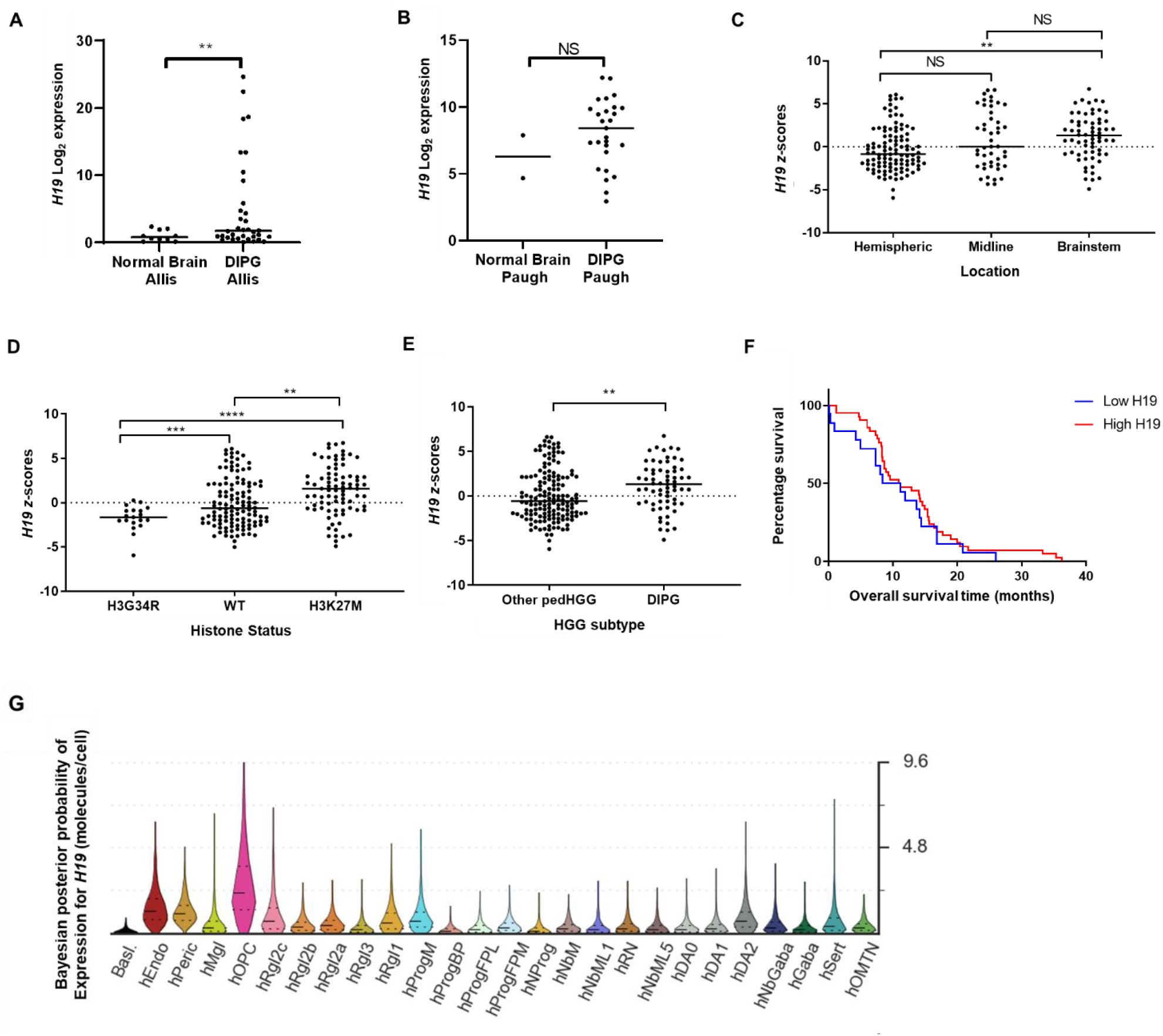
*H19* is associated with DIPG. **(A)** Clinical expression data of *H19* in “Allis - 45 - custom - ilmnht12v4” dataset analysed on R2 platform for normal brain (n=10) and DIPG tissue (n=35) (23,24). **(B)** Clinical expression levels of *H19* in “Paugh - 37 - MAS5.0 - u133p2” dataset analysed on R2 platform for normal brain (n=2) and DIPG tissue (n=27). **(C)** *H19* expression in patient samples with pedHGG located in either hemisphere (n=107), midline (n=46) or brainstem (n=65) analysed with paediatric Cbioportal platform (25). **(D)** *H19* expression in patient samples with pedHGG carrying either H3G34R (n=19), non-mutated/WT (n=118) or H3K27M (n=83) analysed with paediatric Cbioportal platform. **(E)** Expression of *H19* in histologically confirmed DIPG samples (n=65) or other pedHGGs (n=153). **(F)** Kaplan-Meïer curve for overall survival time (months) of DIPG patients with either high or low levels of *H19* (median expression cut-off). **(G)** Violin plot showing Bayesian posterior probability of *H19* expression in different cell types during neurodevelopment. Data in B, C and F were analysed by unpaired two-tailed t-test with Welch’s correction for B and C. Data in D and E were analysed with one-way ANOVA and Dunnet’s post-hoc test for multiple comparison. All Data are shown as mean±SD. Significant values are \*\**P*<0.01, \*\*\**P*<0.001 or \*\*\*\**P*<0.0001.

### H19 is required for DIPG cell viability but not migration

To investigate the functional role of *H19*, expression of this transcript was analysed in a panel of human DIPG cell lines and normal astrocytes. The malignant cell lines showed higher levels of *H19* lncRNA expression, compared to normal astrocytes. VUMC-DIPG-A cells expressed the highest levels of *H19* (Figure 1A.). Biologically, VUMC-DIPG-A cell line bears the H3.3K27M mutation, whereas SU-DIPG-IV cells carry the H3.1K27M mutation (Supplementary Table 1). Cell fractionation assays demonstrated that *H19* lncRNA is ~3-fold enriched in the cytoplasm of both VUMC-DIPG-A and SU-DIPG-IV cells, compared with the nucleus (Figure 2B.)

To understand the cellular function of *H19*, we performed gene silencing studies on VUMC-DIPG-A cells, using locked-nucleic acid (LNA) antisense oligonucleotides. Three LNAs were designed and tested in this cell line. Two days post-transfection, *H19* expression vs. control was 0.47±0.37 with LNA1, 0.16±0.09 with LNA2 (*P*<0.05) and 2.04±2.59 with LNA3 (Figure 2C). LNA2 was therefore selected for cell proliferation assays. We found that cell numbers failed to increase in the presence of LNA2 induced a significant and durable reduction in the number of cells at day 4 and 7 (*P*<0.01 and *P*<0.05, respectively) (Figure 2D). VUMC-DIPG-A cells were also imaged using bright-field microscopy, which showed morphological changes such as detached cells indicative of cell death upon exposure to LNA2 (Figure 2E). We also observed that LNA2 induced a dose-dependent inhibition of DIPG cell numbers, with an estimated IC50 of 52.4±1.4 nM at day 4 measure by trypan blue-exclusion (Figure 2F). In order to determine whether *H19* knock-down attenuated cell proliferation by inducing caspase-dependent apoptosis, caspase 3/7 assays were carried out at day 4. *H19* silencing via LNA2 triggered a significant increase in caspase 3/7 activity (*P*<0.01) (Figure 2G). Notably, *H19* knock-down in non-neoplastic astrocytic cells did not significantly (Figure 2H) To investigate whether *H19* was involved in the migration of VUMC-DIPG-A cells, a wound healing assay was performed 24h post-transfection. *H19* knock-down did not affect the migration of VUMC-DIPG-A cells in this experimental design (Figure 2I.). Overall, these functional assays suggested that *H19* modulates DIPG-H3K27M cell proliferation and apoptosis.

**Figure 2.**
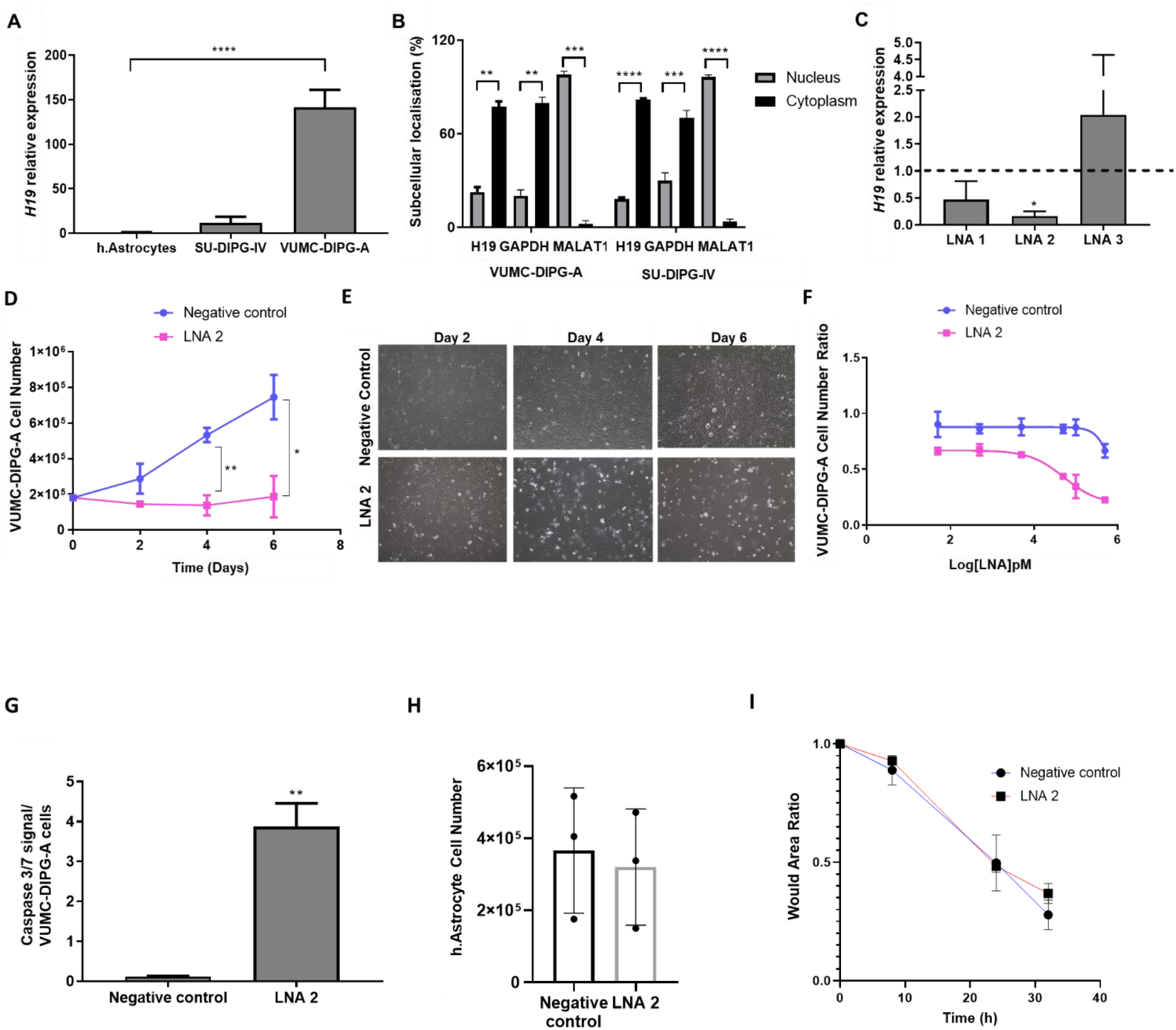
*H19* is required for DIPG cell proliferation but not for migration. **(A)** Expression (qPCR) of *H19* lncRNA in a panel of DIPG cancer cells and a non-neoplastic human astrocyte cell line (n=3). **(B)** Subcellular localisation of *H19* in VUMC-DIPG-A and SU-DIPG-A cells. Internal control included GAPDH (cytoplasm control) and MALAT1 (nuclear control) (n=2 and n=3, respectively). **(C)** *H19* RNA relative expression on LNA-silenced VUMC-DIPG-A cells (LNA1, LNA2 and LNA3) to negative control LNA-silenced VUMC-DIPG-A cells (n=3). **(D)** Trypan blue exclusion cell number of VUMC-DIPG-A after 2, 4 and 6 days’ post-transfection with LNA2 or negative control LNA (n=3). **(E)** Representative brightfield image of LNA2- or negative control LNA-transfected VUMC-DIPG-A cells after 2,4 and 6 days. **(F)** Dose response (0.05 to 500 nM) effect of LNA7-silencing on VUMC-DIPG-A cell number (n=2). **(G)** Cell number of normal human astrocytes after transfection with either LNA2 (50nM) or negative control LNA at day 4 (n=3). **(H)** Cell viability-adjusted activity of caspase 3/7 after 4 days of LNA2-mediated H19 knock-down in VUMC-DIPG-A cells (n=3). **(I)** Wound healing assay showing wound area ratio closing at 8, 24 and 32 h post-scratch (n=3). Data on A was analysed with one-way ANOVA followed by Dunnet’s multiple comparision test. Data on B, C and D were analysed with two-way ANOVA with Sidak’s post-hoc test for multiple comparison. Data are shown as mean±SD. Significant values were \**P*<0.05, \*\**P*<0.01, \*\*\**P*<0.001 or \*\*\*\**P*<0.0001.

### Let-7a-5p affects DIPG-H3K27M cell proliferation

To determine the mechanism of action behind *H19*-driven cell proliferation, we compared the transcriptome of *H19*-knockdown vs control DIPG cells. RNA-sequencing analysis revealed 449 up-regulated and 622 down-regulated mRNAs upon *H19* knockdown using LNA2 in VUMC-DIPG-A cells (Figure 3.A, Supplementary Table 2). Gene set enrichment analysis revealed that the most up-regulated protein-coding genes were associated with: tumour necrosis factor alpha (TNFα) signalling, apoptosis or KRAS signalling. Down-regulated genes were associated to: epithelial and mesenchymal transition, UV response or TGFβ signalling (Figure 3B).

Cytoplasmic lncRNAs often interact with miRNAs. Typically, these lncRNAs bind to complementary miRNAs, thereby displacing the interaction of the miRNA and its mRNAs targets; this results in increase translation of the target mRNAs (26). We therefore decided to test the hypothesis that *H19* exerts its oncogenic function by buffering onco-suppressive miRNAs. As first step, we searched the TARBASE dataset, and identified 11 miRNAs that are validated targets of *H19* (Figure 3C, Supplementary Table 3). Notably, the *let-7* miRNA family was highly represented in this list. Hence, we decided to focus further investigation on *let-7a-5p*, whose direct binding to *H19 has* been demonstrated experimentally (27). However, it is still unknown whether *let-7a-5p* affects DIPG cell proliferation.

To test our hypothesis, we transfected a miRNA mimic of *let-7a-5p* in DIPG cells. Transient overexpression of *let-7a-5p* led to a significant reduced VUMC-DIPG-A cell proliferation 5 days post-transfection (Figure 3D). Morphologically, *let-7a-5p*-overexpressing VUMC-DIPG-A cells showed no obvious differences, compared to those transfected with negative control mimic (Figure 3E). These results suggested *let-7a-5p* is involved in DIPG-H3K27M cell proliferation.

To identify which mRNAs are likely to be targeted by both *H19* and *let-7*, the list of *H19*- down-regulated mRNAs (622) was cross-referenced with the list of *let-7a-5p* predicted mRNAs (1207) extracted from TargetScan 7.0 (Figure 3F). There were 64 overlapping mRNAs, but only 8 of these were positively and significantly (*P*<0.0001) correlated with *H19* expression in clinical samples using data of pedHGG from paediatric Cbioportal. SULF2 and OSMR were the top two protein-coding genes that correlated with *H19* expression (Supplementary Table 4). *Let-7a-5p* had a predicted 6mer and 8mer binding site for *SULF2* and *OSMR*, respectively (Figure 3G). in keeping with our mechanistic model, the expression of *SULF2* (*P*<0.01) and *OSMR* (*P*=0.052) was reduced in *H19*-silenced VUMC-DIPG-A cells (Figure 3H). These results suggest that *H19* potentially sponges *let-7a-5p* thereby causing the up-regulation of *let-7a-5p-target* mRNAs (such as *SULF2* and *OSMR*).

**Figure 3.**
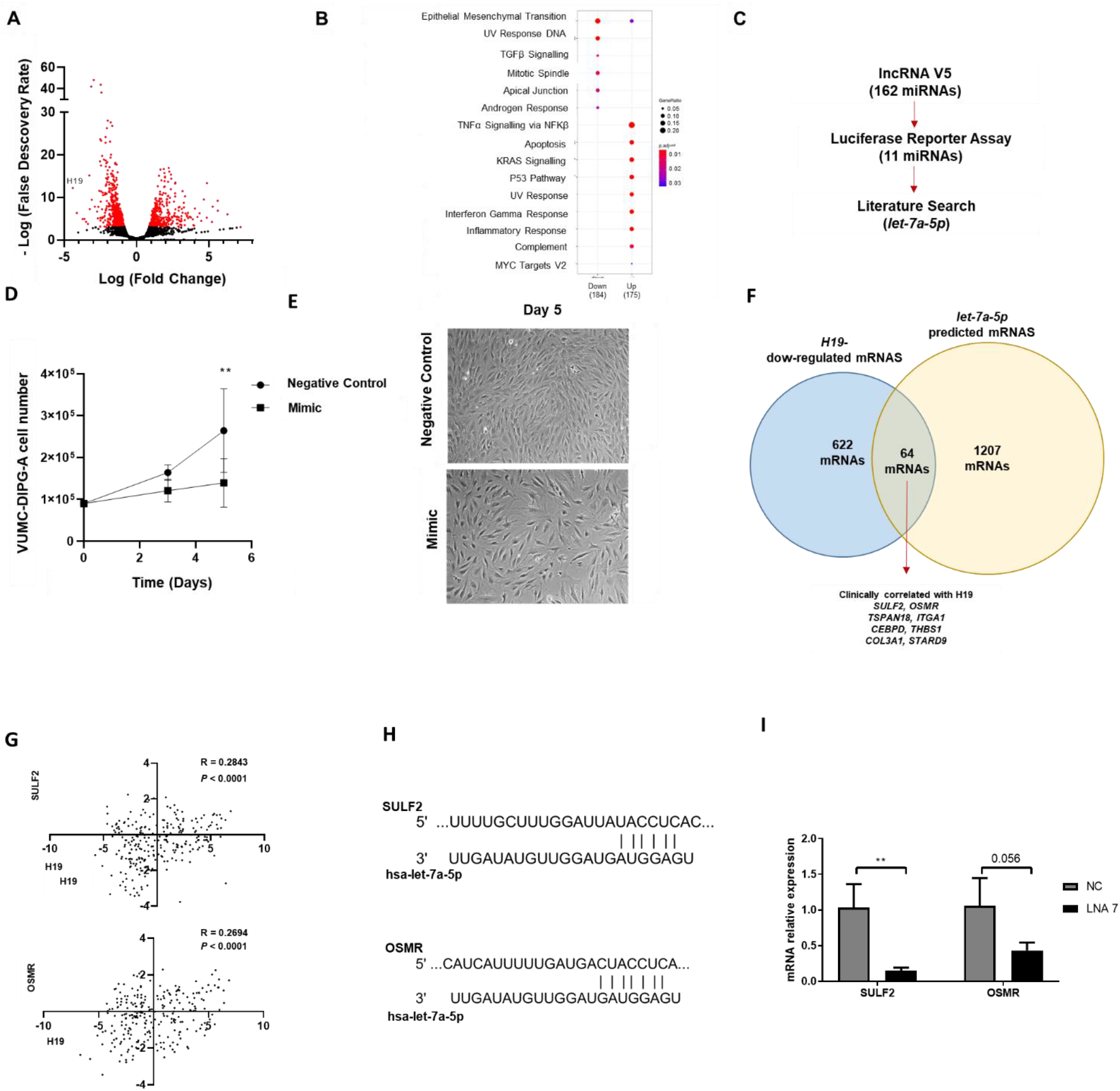
*let-7a-5p* reduces DIPG-H3K27M cell proliferation and migration. **(A)** Volcano plot of RNA-seq expression on *H19*-silenced VUMC-DIPG cells (red dots are *P*<0.01). **(B)** Gene set enrichment analysis of up-regulated and down-regulated (log_2_FC < or > 0.58) mRNAs upon *H19* silencing in VUMC-DIPG-A cells. **(C)** Process diagram for identification of *H19*-previously validated miRNA targets with lncRNA v5 database and luciferase binding assay. **(D)** Trypan blue-exclusion cell number on *let-7a-5p*-overexpressing VUMC-DIPG-A cells at day 3 and 5 post-transfection. **(E)** Brightfield images of *let-7a-5p* mimic and control (scramble oligonucleotide) overexpressing VUMC-DIPG-A cells (n=4). **(F)** Venn diagram of cross-referencing *H19*-silenced mRNAs with *let-7a-5p* predicted mRNA targets. Overlapping mRNAs were analysed for clinical correlation with *H19* in paediatric Cbioportal. **(G)** Correlation analysis of *H19* expression and SULF2 or OSMR in ICR 2017 pedHGG dataset in paediatric Cbioportal platform (25). **(H)** Graphical representation of predicted binding site between *let-7a-5p* and SULF2 (top) or OSMR (bottom) mRNA. **(I)** SULF2 and OSMR expression in *H19*-silenced VUMC-DIPG-A cells 2 days after transfection analysed by qPCR (n=3). Data on B were analysed with two-way ANOVA and Sidak’s post-hoc for multiple comparison. Data on I were analysed with unpaired two-tailed t test. Data on G was analysed using Spearman correlation. Data are shown as mean±SD and \*\**P*<0.01.

## DISCUSSION

In this study, we showed that *H19* is up-regulated in DIPG, and that silencing *H19* decreased DIPG-H2K27M cell proliferation and increased caspase 3/7 activity. To the best of our knowledge, this is the first functional characterisation of a lncRNA in DIPG. The oncogenic role of *H19* has been previously proposed for a wide range of malignancies including adult high grade glioma (15). In keeping with our findings, several lines of evidence showed *H19* silencing led to decreased cell proliferation in other malignancies; interestingly, this effect was explained by a wide range of mechanisms of action, which appear to be at least in part cell-specific (28–30).

The origin of DIPG is controversial, but recent research suggests that these malignancies originate from the partial differentiation of neural stem cells into oligodendrocyte precursor (OPC)-like cancer cells (4,31). Here we show that *H19* is up-regulated in OPC cells, during brain development. In keeping with this finding, it has been shown that *H19* is down-regulated upon full differentiation of OPCs into oligodendrocytes (32). Taken together, our results and previous publications suggest that *H19* may be implicated in the proliferation of OPC-like cancer cells, which are the cells of origin in DIPG.

Mechanistically, *H19* is a cytoplasmic lncRNA that can sponge miRNAs to modulate gene expression (26). Several miRNAs have been shown to directly interact with *H19*, including *miR-141-3p, miR-200b-3p* and *let-7a* (33–35). It is therefore likely that the effects of *H19* on normal and cancer cells are determined by the complex interplay of a pool of cell specific miRNAs expressed in each cell type. To discover the miRNAs that mediate *H19* function in DIPG, we crossed the list of *H19* targets with the list of miRNAs able to bind to the mRNAs down-regulated upon *H19* silencing. Based on this analysis, we decided to further investigate *let-7a-5p*, because this miRNA had been shown to interact with *H19* in other contexts (35–37), because it had been shown to down-regulate 64 *H19*-dependent mRNAs.

In keeping with our hypothesis, overexpression of *let-7a-5p* led to the reduction of DIPG cell proliferation; this observation suggested that *H19* buffers the onco-suppressive *let-7a-5p*, thereby increasing DIPG cell proliferation. However, *let-7a-5p* overexpression was not as cytotoxic as that of *H19* silencing; this observation suggests *let-7a-5p* is only partially responsible for *H19* oncogenic function.

Interestingly, SULF2 increases cell proliferation, invasion, mobility and adhesion in MCF-7 and MDA-MB-231 breast cancer cell lines (38). OSMR promotes proliferation, metastasis and EMT in prostate cancer (39) OSMR also increases mitochondrial respiration and radio-resistance in glioblastoma stem cells (40).

Cell proliferation is defined as the balance between cell survival and cell death. Our results indicate that *H19* is able to drive cell survival and suppress cell death in DIPG cells. However, *let-7a-5p* seems only to affect cell growth. Therefore, it seems likely that other miRNAs mediate the anti-apoptotic function on *H19* in DIPG.

The primary aim of this work was to functionally characterize a DIPG-specific lncRNA, and to identify therapeutic targets for this incurable malignancy. For this reason, silencing experiments were carried out with LNAs instead of more conventional siRNAs. LNAs are more stable in biological media and are currently being investigated in clinical trials for the treatment of neurological disorders and malignancies (41–44). Here, we report that an *H19*- targeting LNA (LNA2) had an *in vitro* IC50 that was lower than typical maximum plasma concentrations of LNAs in clinical trials (2000-6000 ng/ml) (45). Future *in vivo* studies are warranted to establish whether LNA2 is effective in clinically relevant animal models of DIPG-H3K27M.

In conclusion, this work has shown for the first time the role of an oncogenic lncRNA in DIPG, which is likely to control multiple pathways in DIPG-H3K27M. We hope this work will encourage DIPG researchers to further explore the function of lncRNAs and to test their role as therapeutic agents.

## MATERIAL AND METHODS

### R2 platform analysis

*H19* lncRNA was queried in R2: Genomics Analysis and Visualization Platform (https://hgserver1.amc.nl/cgi-bin/r2/main.cgi) to assess clinical relevance in DIPG tissue and normal brain tissue. The datasets used for this analysis were GSE26576 (R2 id was ps_avgpres_gse26576geo37_u133p2 and it was referred as “Paugh”) and GSE50021 (R2 id was ps_avgpres_gse50021geo45_ilmnht12v4 and it was referred as “Allis”) (23,24). Data was sorted based on “disease_type” (Paugh) and “cell_type” (Allis). Only DIPG and normal brain tissue were compared.

### Paediatric Cbioportal analysis

The paediatric Cbioportal analysis was carried out with *H19* lncRNA (https://pedcbioportal.kidsfirstdrc.org/) to assess the difference of expression of *H19* in paediatric high grade gliomas. Data was obtained from publicly available Institute of Cancer Research (ICR) dataset from 2017 (25) and sorted based on either 1) type of histone modification (wild type, WT; H3.1K27M and H3.3K27M, grouped as H3K27M; H3G34R), 2) brain location (hemispheric, midline or brainstem) or 3) histological classification (DIPG or other paediatric high grade glioma, which included anaplastic glioma and glioblastomas). Data was also used to perform Kaplan-meïer analysis sorted based on median expression of H19 in Graphpad Prism 7 (Graphpad Software).

### Single cell analysis

Open-access dataset on single-cell analysis of midbrain in development (http://linnarssonlab.org/ventralmidbrain/). *H19* was queried and expression in human cells was analysed.

### Cell culture

Primary human foetal cortical normal astrocytes were a gift of Prof. DK Male (The Open University) and were cultured as described elsewhere (46). The VUMC-DIPG-A (H3.3 K27M) cell line were kindly provided by Dr.Esther Hulleman (VUMC Cancer Center, Amsterdam, The Netherlands) (22). SU-DIPG-IV (H3.1 K27M) cells were kindly provided by Dr. Michelle Monje (Stanford University, California, USA). (47). VUMC-DIPG-A cells were cultured in 1:1 DMEM-F12 and Neurobasal-A cell media and supplemented with 1% (v/v) glutamax supplement, 1% (v/v) antibiotic-antimycotic solution, 10mM HEPES, 1% (v/v) MEM non-essential amino acid solution and 1mM sodium pyruvate (Thermofisher), hereafter referred as TBM media. VUMC-DIPG-A media was also supplemented with 10% heat-inactivated foetal bovine serum (Merk). SU-DIPG-IV were cultured in TBM media supplemented with 2% (v/v) B27 supplement without vitamin A (Thermofisher), 20 ng/mL bGFG, 20 ng/mL.EGF, 10 ng/mL PDGF-AA, 10 ng/mL PDGF-BB (Peprotech) and 5IE/mL Heparin (Merk). Cells were grown in adherence and passaged using 0.25% (v/v) trypsin-EDTA (Merk). All cells were cultured at 37 °C in a humidified environment containing 5% CO_2_.

### Analysis of gene expression

RNA was isolated from cells in culture using the RNeasy plus mini Kit (Qiagen) according to manufacturer’s instructions. Reverse transcription was carried out using 1 μg RNA per reaction using the high capacity cDNA reverse transcription kit (Applied Biosystems) according to kit instructions. For RT-qPCR analysis, 10 ng of cDNA were loaded in duplicated per sample using Taqman Universal mastermix and primers. Taqman primers used in this study included: *H19*, (Hs00399294_g1); GADPH (Hs02786624_g1), SULF2, (Hs01016480_m1); OSMR (Hs00384276_m1).

### Subcellular localization of *H19*

RNA was isolated from cells in culture in nuclear and cytoplasmic fractions which were separated using the PARIS kit (Ambion) according to manufacturer’s instruction. For validation and localisation studies RT-qPCR was used with the probes MALAT1 (Hs00273907_s1), GAPDH (Hs02786624_g1) and HPRT1 (Hs02800695_m1)

### LNA reverse-transfection

Knock-down studies were performed using the reverse transfection method. Cells were seeded in a 6-well (180,000 cells/well) or 96-well plate (5,000 cells/well) with lipid:LNA mixture prepared using Oligofectamine (Invitrogen) as per manufacturer’s protocol. Final LNA concentrations were 100 nM. All LNAs were designed using Qiagen LNA design tool and targeting sequences were ACTAAATGAATTGCGG (LNA 1), AATTCAGAAGGGACGG (LNA 2) and GACTTAGTGCAAATTA (LNA 3). Control LNA were also purchased from Qiagen.

### Cell proliferation assay

LNA-transfected cells were seeded on 6-well plates and grown at specific times as specified in figure legends. On the day of analysis, cells were trypsinized, centrifuged and resuspended in HBSS for cell counting using trypan blue (Merk) exclusion method with either a cell haemocytometer or an automatic cell counter (LUNA, Logos Bioscience).

### Caspase 3/7 assays

Cells were plated in a white, flat-bottomed 96-well plates and treated with 100nM of LNAs as previously described. On day 4 post-transfection, Caspase-Glo reagent was added per well (Promega) according to manufacturer’s instruction was added to well and incubated for 1.5 h. Luminescence was then quantified using the BMG polarSTAR plate reader (BMG Labtech).

### Wound Healing assay (migration)

VUMC-DIPG-A cells were reverse-transfected for 18h with 50 nM LNA2 as these did not yield a significant reduction in viable cells as determined by cell proliferation assays. After 18 h, a scratch was made in the cell monolayer using a sterile P20 pipette tip. Cells were imaged periodically across the wound until the wound was closed. Images were analysed using the ImageJ.

### RNA sequencing analysis

VUMC-DIPG-A cells were reverse-transfected with 100 nM of LNA and RNA isolated 48h post-transfection. NGS libraries were prepared according to the Ion AmpliSeq Library Kit Plus user guide (Thermo Fisher Scientific). For each sample, 100 ng of total RNA was reverse transcribed using the SuperScript VILO cDNA synthesis Kit (Thermo Fisher Scientific). The resulting cDNA was then amplified for 10 cycles by adding PCR Master Mix and the AmpliSeq Human Transcriptome Gene Expression Kit panel, targeting over 20,000 genes (> 95% of the RefSeq gene database). Amplicons were digested with the proprietary FuPa enzyme to generate compatible ends for barcoded adapters ligation. The resulting libraries were purified using AmpureXP beads (Agencourt) at a bead to sample ratio of 1.5× and eluted in 50 μL low TE buffer. Libraries were then diluted 1:10000 and quantified by qPCR using the Ion Universal Quantitation Kit (Thermo Fisher Scientific). Individual libraries were diluted to a 50 pM concentration, combined in batches of 8 libraries, loaded on an Ion 540™ chip using the Ion ChefTM instrument and sequenced on an Ion S5™ instrument (Thermo Fisher Scientific). QC was manually performed for each sample based on the following metrics; number of reads per sample >8*10^6^, valid reads > 90%. Raw data was processed automatically on the Torrent ServerTM and aligned to the reference hg19 AmpliSeq Transcriptome fasta reference.

The AmpliSeqRNA plugin was used to determine valid matches to amplicon target regions in the panel using the hg19 AmpliSeq Transcriptome bed file. Each sample had more than 5*10^6 total aligned reads. At least 10 mapped reads per amplicon in more than 3 samples were used to filter out lowly covered amplicons. Two analyses pipelines were used separately. Log-fold changes reported by DESeq2 (10.1186/s13059-014-0550-8) were shrunk with apeglm (10.1093/bioinformatics/bty895) available within DESeq2 Bioconductor package(package version: 1.30.1, 10.18129/B9.bioc.DESeq2). Genes were reported as significant with log-fold change > 0.58 and svalue < 0.01. Analysis were performed in R (version: 4.0.3, R Core Team (2020) using RStudio interface (version 1.3.1073).

Biological function gene ontology was analysed with Gene Set Enrichment Analysis (http://www.gsea-msigdb.org/gsea/msigdb/index.jsp) and “Hallmark” feature using genes grouped as up-regulated (log2FC>0.58 and svalue<0.01) or down-regulated (log_2_FC<-0.58, svalue<0.01).

### miRNA mimic studies

VUMC-DIPG-A cells were reverse transfected with 30 nM negative control (NC, scramble control) or *let-7a-5p* mimic using the method previously described. The sequences used were obtained from Thermo Fisher and were as follows: *let-7a-5p* mimic (MC31809) and NC mimic (4464058).

### Analysis of miRNA expression

For miRNA analysis, RNA was isolated using miRNeasy (Qiagen) according to manufacturer’s instructions. Reverse transcription was carried out using the miRNA cDNA systhesis kit (Applied biosystems) according to kit instructions using the reverse transcription primers provided with *let-7a-5p* (000377) or *U6* snRNA (001973) using 10 ng RNA per reaction. Then, 1.7 ng of resulting cDNA was used for qPCR analysis using the qPCR taqman probes provided in the kits described for *let-7a-5p* or U6 according to manufacturer’s instructions

### *In silico* prediction of miRNA target genes

Identification of experimentally validated miRNA-lncRNA binding site was carried out with the online openaccess lncBase v.3 database (http://carolina.imis.athenainnovation.gr/diana_tools/web/index.php?r=site%2Ftools) (48). Only miRNAs whose binding has been previously demonstrated by luciferase assay were taken into account. Prediction for miRNA-mRNA interactions was investigated using open-access TargetScan database (http://www.targetscan.org/vert_72/). Predicted target mRNAs were overlapped with mRNAs down-regulated upon *H19* knockdown (RNA-seq) in VUMC-DIPG-A cells. Top mRNAs that were positively correlated (Spearman values < 0.05) with *H19* expression in pedHGG clinical dataset (paediatric Cbioportal, ICR 2017 pedHGG study) were prioritized for validation.

### Statistical analysis

All data are presented as mean ± SD (standard deviation). The number of independent experiments (n) with replicates specified in each legend. Normality was assessed with Shapiro-Wilk test, *P*=0.05. Means were compared using unpaired or paired two-tailed t-test for single comparison and one-way or two-way ANOVA for multiple comparisons. ANOVA analysis were followed by post hoc analysis. Specific post-hoc analysis is specified in each figure legend. All tests were performed using the statistical software GraphPad Prism 7 (GraphPad Software, San Diego, CA, USA). A *P*<0.05 was considered to be statistically significant.

## Supplementary material

**Supplementary Table 1. Clinical characteristics of DIPG cell models.** Table summarizing sex, age and histone 3 gene modification of diffuse midline glioma cell lines (SU-DIPG-IV and VUMC-DIPG-A).

**Supplementary Table 2. List of de-regulated genes upon *H19* silencing in VUMC-DIPG-A cells.** Cells were silenced with 100 nM of *H19*-targetting LNA2 and gene expression was analysed by RNA-sequencing. Results show genes with s.value <0.01 and log2 fold change (lfcSE) >0.58.

**Supplementary Table 3. List of miRNAs that are experimentally validated to be targeted by *H19* lncRNA.** LncBase v.2 analysis of miRNA that have been previously reported positive luciferase reporter assay of binding with *H19* lncRNA.

**Supplementary Table 4. List of correlated *let-7* target mRNA and *H19* lncRNA.** Analysis was carried using clinical data from paediatric gliomas dataset (ICR 2017) in paediatric cbioportal.

## Acknowledgements

Special thank you to Amanda Mifsud for supporting this project via Abbie’s Army funding. We would like to thank Perla Pucci, Maryam Latarni, Mark Hirst and Sushila Rigas for providing feedback and advice for the project.

