## Supplementary material for "The long non-coding RNA *H19* drives the proliferation of diffuse intrinsic pontine glioma with H3K27 mutation": Supplementary Table 1 (1).pptx

### Slide 1
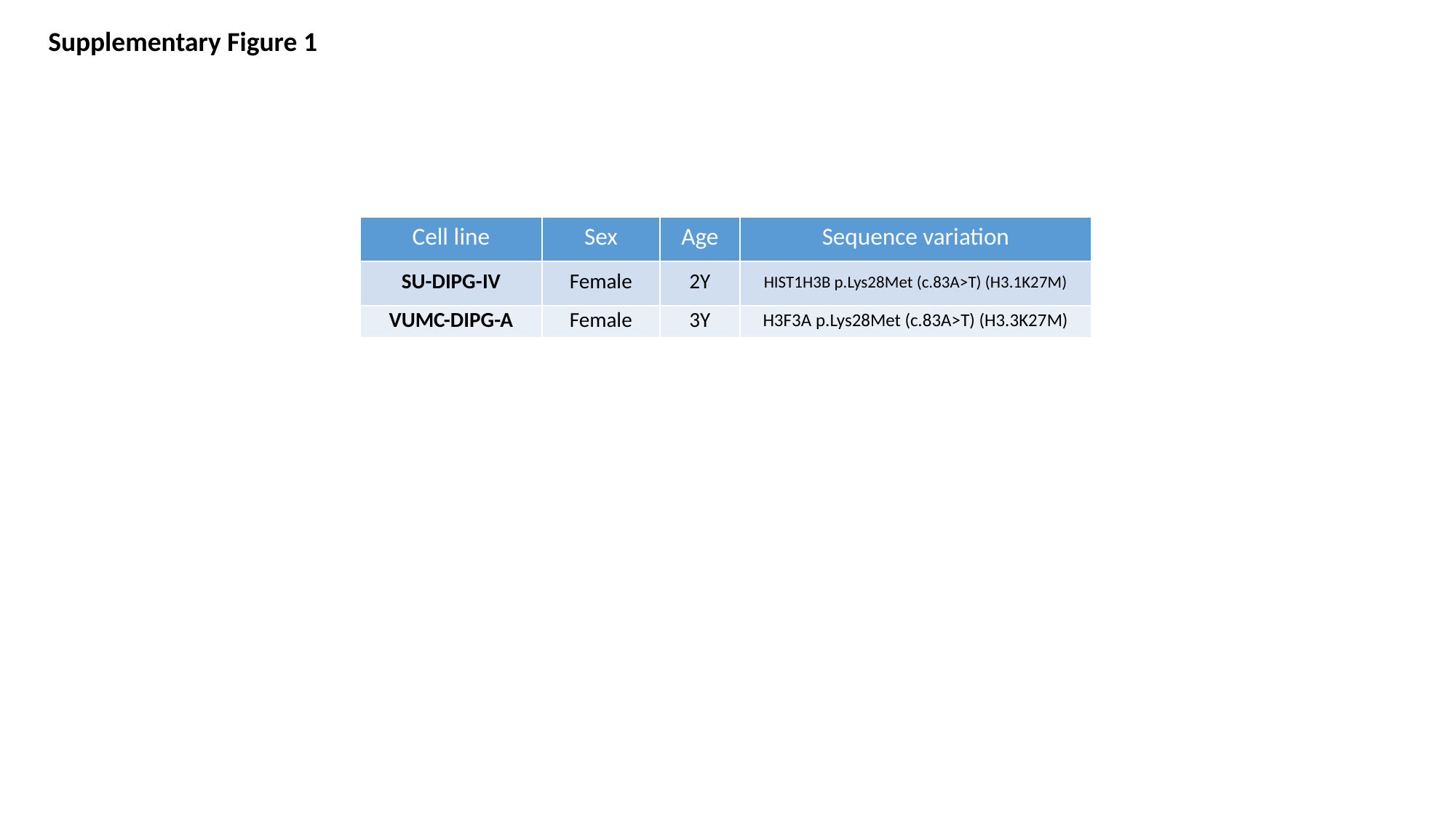

Supplementary Figure 1
| Cell line | Sex | Age | Sequence variation |
| --- | --- | --- | --- |
| SU-DIPG-IV | Female | 2Y | HIST1H3B p.Lys28Met (c.83A>T) (H3.1K27M) |
| VUMC-DIPG-A | Female | 3Y | H3F3A p.Lys28Met (c.83A>T) (H3.3K27M) |
